# Computational screening and automatic filtering for the discovery of novel inhibitors of TMPRSS2, a type II transmembrane serine protease

**DOI:** 10.1101/2025.09.09.675039

**Authors:** Luca Tongiorgi, Simone Albani, Laura Rigobello, Cristina Maccallini, Francesco Musiani

## Abstract

Transmembrane Serine Protease 2 (TMPRSS2) is a membrane protein of the type II serine protease family of enzymes implied in epithelial homeostasis. It is involved in several diseases, notably prostate cancer and SARS-CoV-2 infections. Over the years, only a few tested TMPRSS2 inhibitors showed consistent results. This prompted us to select it as target of structure-based virtual screening, to search for novel inhibitors among a library of 475,770 small molecules. Two sets of TMPRSS2 structures were selected, one taken from molecular dynamics simulations, the other from recently solved X-ray crystallographic structures. We designed a workflow to filter docking results in a reproducible way, allowing for a faster and more reliable selection. The program uses four metrics: the pose consistency of the ligand, docking score, number of interactions with key protein residues, and cluster analysis. This led to the selection and visual inspection of two sets of 500 compounds, which yielded 10 reasonable hit candidates.

## Introduction

### TMPRSS2 in Human Physiology and Disease

Transmembrane Serine Protease 2 (TMPRSS2, EC:3.4.21.122) is an important member of the type II Transmembrane Serine Protease (TTSP) family, belonging to the hepsin subgroup.^1^ These proteins are involved in various proteolytic cascades and mediate important physiological functions such as digestion,^2^ hearing,^3^ liver metabolism,^4^ blood pressure regulation,^5^ degradative remodelling of the extracellular matrix and various other key epithelial homeostasis roles.^6^ While clear evidence regarding TMPRSS2 physiological roles remains scarce,^7,8^ its involvement in a number of pathological processes has instead been described in greater detail.

TMPRSS2 is highly expressed in prostate epithelial cells and was found to be dysregulated in prostate cancer cells.^9^ The protein overexpression leads to looser extracellular environment and thus facilitates the migration of cancerous cells.^10^ Moreover, the TMPRSS2 gene expression is under the control of androgen receptor stimulation, due to the presence of an Androgen Responsive Element (ARE) in its 5’ untranslated region, a relevant feature since androgen signalling plays a major role in the disease progression.^11^ TMPRSS2 is also highly expressed in human respiratory tissues, and its role in the cell entry of enveloped respiratory viruses has been thoroughly described for influenza viruses and coronaviruses, as it cleaves and activates viral surface proteins to facilitate membrane fusion and entry into host cells.^12–18^

### Structure and catalytic function

The most studied isoform is composed of 492 residues and its structure can be divided into a short N-terminal intracellular domain, a type II transmembrane domain (i.e., composed of a single α-helix spanning the cell membrane), an ectodomain. The ectodomain includes the so-called stem region, composed of a Low-Density Lipoprotein Receptor Class A (LDLRA) domain, a Scavenger Receptor Cysteine-Rich (SRCR) domain, and finally the C-terminal Serine Protease (SP) domain (**Fig. 1a**),^1^ which harbours the conserved catalytic triad, comprising Ser441, His296 and Asp345 (**Fig. 1b**). Within the active site, various subsites can be identified according to the nomenclature introduced by Schechter and Berger,^19^ among which the S1 subsite is the most important for the protein function. S1 is composed of Asp435, Ser436, Gly462, Gly464, and Gly472 residues, which form a deep cavity to accommodate an arginine or lysine corresponding to the P1 substrate site. The protease specificity towards monobasic amino acids is largely influenced by the negatively charged Asp435 residue, which forms a fundamental salt bridge with the positively charged basic function of the substrate and is thus termed the “substrate recognition” site.^20^ The so-called “oxyanion hole” is another relevant subsite, formed by the secondary amide NHs of Gly439 and Ser441 backbone, which contribute to activation and stabilisation of the oxyanion resulting from the formation of the tetrahedral complex.^21^

**Fig. 1.**
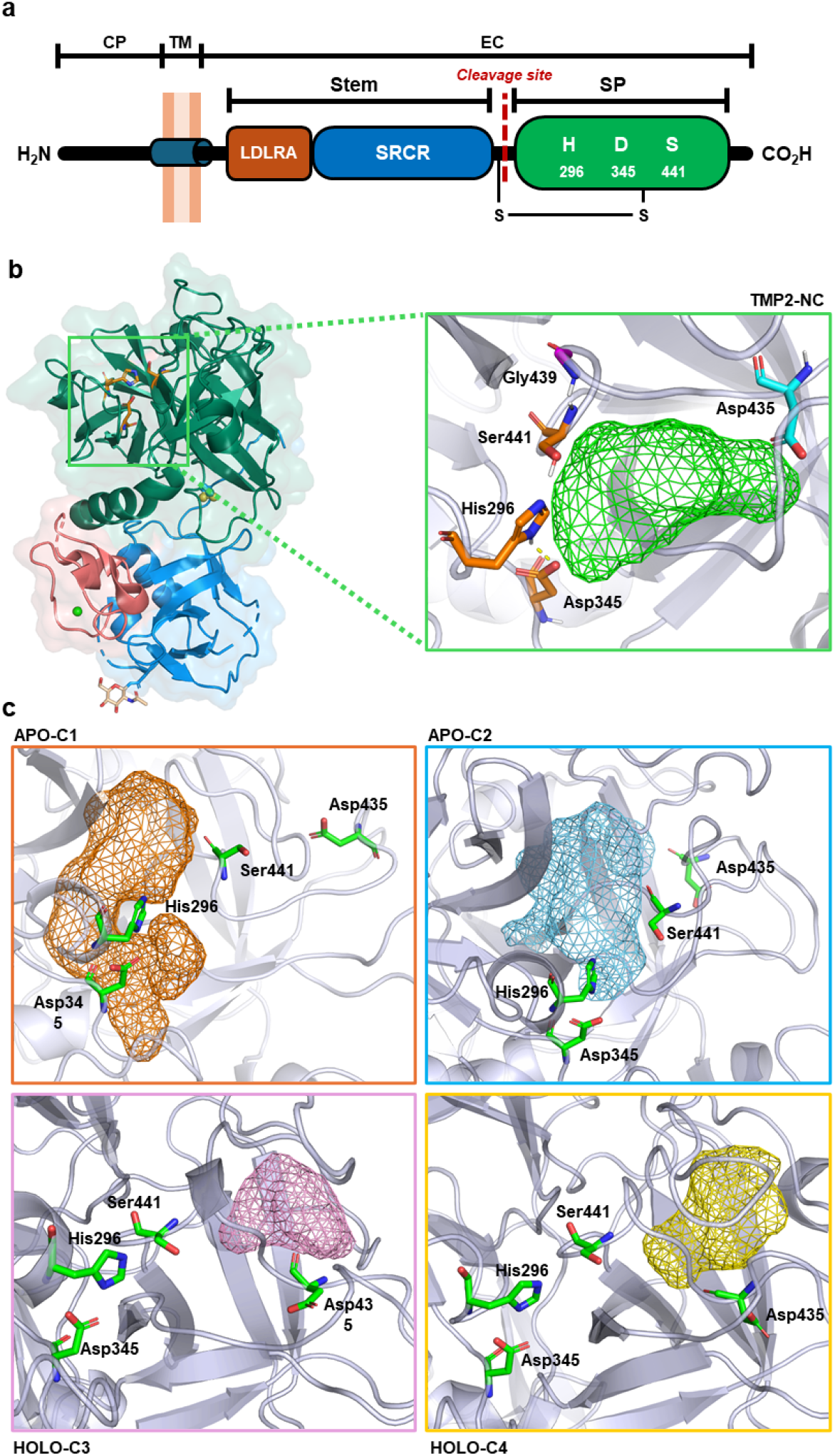
(a) Scheme of TMPRSS2 molecular structure. (b) X-ray crystal structure of the ectodomain of TMPRSS2 in cartoon representation (PDB ID: 7Y0F). The LDLRA is represented in red, with a Calcium ion in sphere representation colored green. The SRCR is in blue, with the glycosylation site in beige-colored sticks representation. The serine-protease domain is in dark green, with the catalytic triad Ser441, His296 and Asp345 in orange-colored sticks representation. Zoomed in is a detail of the catalytic site of TMP2-NC (PDB ID: 7Y0F), representative for the catalytic sites of the other X-ray crystallographic structures. The volume of the binding site cavity is shown as green mesh. Key residues are shown in sticks representation; the catalytic triad carbon atoms are shown in orange, Asp435 (substrate recognition) carbons are light blue, while Gly439 (oxyanion hole) carbons are magenta. Other non-hydrogen atoms of the key residues are colored according to the standard color scheme. (c) Detail of the binding pockets on the four MD frames. The catalytic triad and Asp435 are shown in stick representation, with carbon atoms green and other non-hydrogen atoms colored according to the standard color scheme. The volume of the pocket for apo-C1 is represented as orange mesh, apo-C2 pocket is light blue and noticeably smaller in size. Holo-C3 and holo-C4 have the volume of an allosteric pocket bordered by Asp435 outlined in pink and yellow respectively, shallower in holo-C3 compared to holo-C4.

In 2022, Fraser *et al*. reported the first X-ray crystallographic structure of human TMPRSS2 in complex with the covalent inhibitor nafamostat (PDB ID: 7MEQ, resolution 1.95 Å), with the LDLRA domain not included in the resolved structure.^6^ In July 2023, Frumenzio *et al*. refined the structure using AlphaFold2, enabling complete modelling of its ectodomain and transmembrane region. They performed molecular dynamics (MD) simulations on both the ligand-free (apo) and ligand-bound (holo) forms identifying four representative conformations, each characterized by a distinct conformational state of the active site, hereafter referred to as apo-C1, apo-C2, holo-C3, and holo-C4 (**Fig.1c**).^22^ Later that year, Wang *et al*. disclosed the first X-ray crystallographic structures of human TMPRSS2 including the LDLRA domain, in complex with various inhibitors (**Fig. 2a**): covalent inhibitors nafamostat (PDB ID: 7XYD, resolution 2.58 Å) (**Fig. 2b**) and camostat (PDB ID: 7Y0E, resolution 2.39 Å), a non-covalent inhibitor UK-371804 (PDB ID: 7Y0F, resolution 2.6 Å) (**Fig. 2b**), and the nafamostat-inspired dual inhibitor 212-148 (PDB ID: 8HD8, resolution 2.4 Å),^23^ respectively referred to as TMP2-nafamostat, TMP2-camostat, TMP2-NC (non-covalent) and TMP2-DI (dual inhibitor) hereafter.

**Fig. 2.**
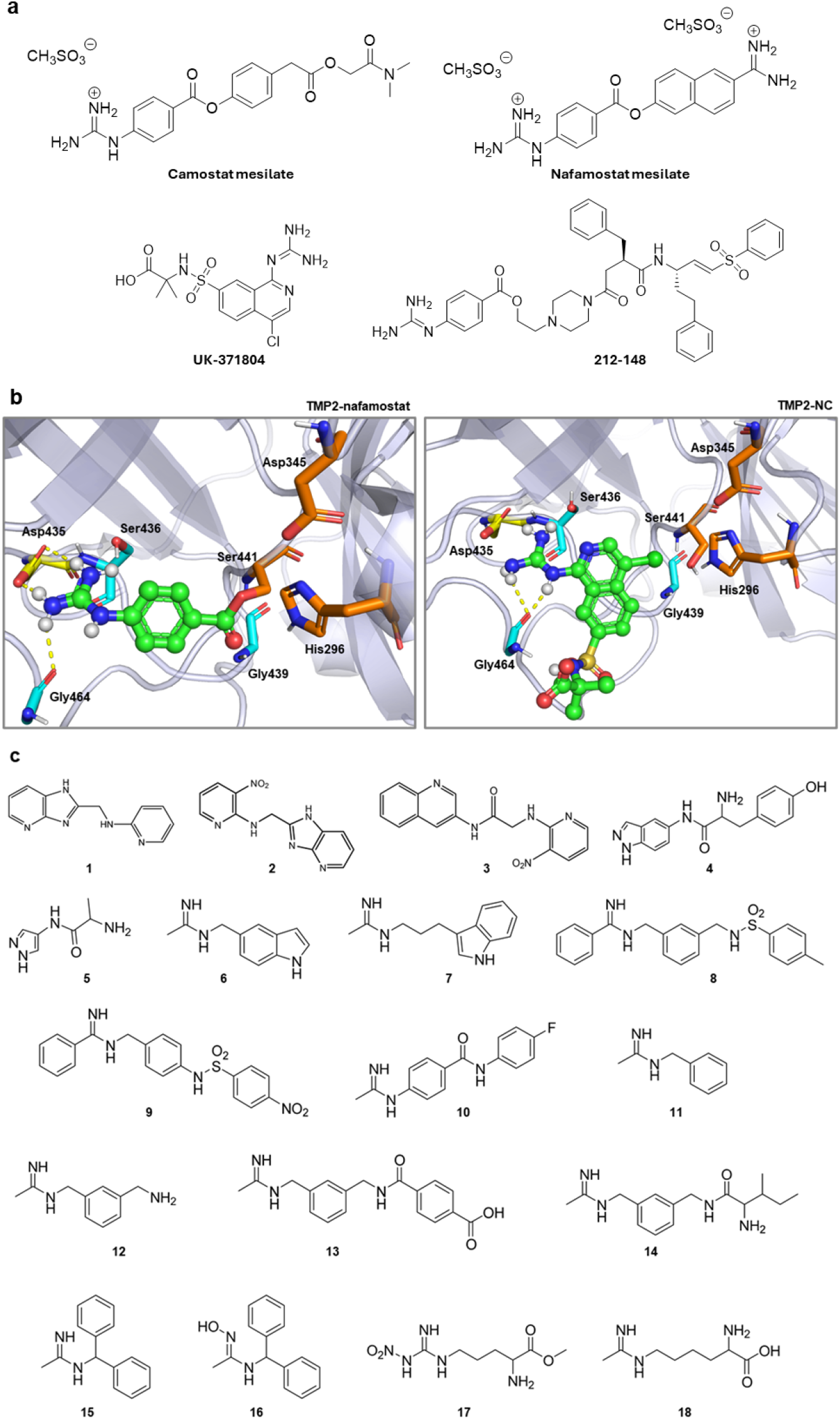
(a) Molecular structures of TMPRSS2 inhibitors camostat mesilate, nafamostat mesilate, UK-371804 and 212-148. (b) Detail of TMP2-nafamostat native complex with the phenylguanidino acyl moiety of nafamostat (left), covalently bound to Ser441 (PDB ID: 7XYD), and detail of the native complex of TMP2-NC with non-covalent inhibitor UK-371804 (PDB ID: 7Y0F) (left). The ligand is shown in ball-and-stick representation with green carbons, while key residues of the catalytic site are shown in sticks. The catalytic triad is shown with orange carbons, Asp435 in yellow, the other interacting residues Ser436, Gly439, Gly464 in light blue. Other non-hydrogen atoms are colored according to the standard color scheme. Interactions between the ligand and the residues are shown as yellow dashed lines. (c) Molecular structures of the 18 initial molecules used as queries to generate the screened library.

### Drug Discovery Efforts Against TMPRSS2

The COVID-19 pandemic spurred intense research into TMPRSS2 inhibitors due to the enzyme’s role in viral entry. Although several compounds showed good *in vitro* or *in silico* results, clinical testing has often been disappointing, partly because of TMPRSS2’s high homology with other proteases.^24^ Camostat mesilate and nafamostat mesilate, both approved in Japan for pancreatitis treatment,^25^ have demonstrated strong TMPRSS2 inhibition and antiviral activity in lung cells against SARS-CoV and MERS-CoV.^26^ Despite encouraging early data and multiple clinical trials, recent meta-analyses report that the clinical outcomes are inconclusive or not significantly better than placebo.^27,28^

In the study that presented the most recent TMPRSS2 X-ray crystal structures, Wang *et al*. also developed a FRET-based assay to evaluate inhibitor activity. The non-covalent inhibitor UK-371804 showed a promising IC50 of 4.59 nM but limited *in vitro* inhibition of the SARS-CoV-2 Omicron variant, and moderate efficacy against the Delta variant with an EC50 of 424.6 nM.^23^ Additionally, the authors designed a bispecific inhibitor, 212-148, targeting both TMPRSS2, and cathepsins L and B, which inhibited TMPRSS2 with an EC50 of 1.38 µM and successfully blocked infection by the SARS-CoV-2 Delta variant.^23^

The search for TMPRSS2 inhibitors has also been strongly supported by computational methods, which address the high cost and long timelines of traditional drug discovery.^25^ Structure-based virtual screening, in particular, has become a key tool for hit discovery, employing molecular docking to enable a rapid evaluation of millions of drug candidates against a target protein by predicting their binding orientations and affinities.^29^ However, docking relies on scoring functions that largely approximate the energy of molecular interactions and assumes a rigid protein structure, often resulting in inaccurate rankings and false positives.^30^ It is for this reason that a scrupulous post-processing should always be conducted on docking results.^30,31^

To improve reliability, post-processing steps like visual inspection are commonly applied, allowing researchers to manually select promising binding modes. While effective, this process is subjective, time-consuming, and often lacks standardised criteria. Techniques like interaction fingerprints, ensemble docking, and cluster analysis help refine results by enhancing data richness, sampling multiple protein conformations, or promoting chemical diversity. Nevertheless, even with the use of these tools visual inspection remains an essential final step in most virtual screening workflows, as it consistently enhances the quality and accuracy of hit selection.^30^

Many of the computational studies on TMPRSS2 were conducted before the publication of its first X-ray crystallographic structure in 2022, and had to rely on homology modeling.^25^ For this reason, the recent publication in 2023 of the most complete structures yet of the protease^23^ could provide new insight and an increased level of accuracy for *in silico* experiments. Our interest was further backed up by the need for novel inhibitors of TMPRSS2, coupled with the relevance of the diseases that this protein is involved in, which altogether prompted us to select it as the target of a structure-based virtual screening study.

### Inducible Nitric Oxide Syntase

Together with TMPRSS2, our interest was placed also on the inducible nitric oxide synthase (iNOS), an enzyme that catalyzes the production of elevated levels of nitric oxide (NO) in response to pro-inflammatory stimuli.^32^ NO is an important pro-inflammatory mediator involved in immune regulation, host defense, neurotransmission, and vascular function.^33^ However, its action can be both beneficial, providing protection against pathogens, or detrimental, damaging healthy tissues.^32^ Indeed, NO and iNOS in particular have been implicated in the pathogenesis of multiple inflammatory disorders.^34^ This study started from known iNOS inhibitors, with the aim of achieving a multi-target effect, by blocking TMPRSS2-mediated viral entry and iNOS-mediated inflammatory cascades synergistically, possibly mitigating the process known as cytokine-storm, typical of SARS-CoV-2 infections.^35^

## Results

The virtual screening campaign began with a set of 18 compounds originally developed as iNOS inhibitors.^36–38^ These molecules were selected for their ionizable nitrogen-rich groups, which we hypothesized could support binding modes similar to those of known TMPRSS2 inhibitors (**Fig. 2c**). In particular, the most common moiety is a terminal acetamidine group, which shares a high stereo-electronic similarity with the guanidine group of several known inhibitors, e.g. camostat, nafamostat and UK-371804. In order to obtain a larger library, Smallworld search engine^39^ was used to perform a similarity search in the Enamine REAL database (version 22Q1),^40^ using default search parameters and the 18 initial compounds as queries. This comprehensively yielded a first pool of 838,646 molecules (**Fig. 3a**). Duplicates were removed based on both SMILES and manufacturer ID, resulting in a library of 271,161 unique purchasable drug-like compounds. Next, the most probable protonation states at pH 7.4 were calculated, and molecules with unassigned stereochemistry had all possible R/S and E/Z configurations enumerated by generating the corresponding stereoisomers. The result was a library of 480,688 compounds. In this study we performed ensemble docking on the two separate sets of receptors (MD frames apo-C1, apo-C2, holo-C3, holo-C4, and X-ray crystallographic structures TMP2-nafamostat, TMP2-camostat, TMP2-NC, TMP2-DI), defining binding sites around catalytic Ser441. In holo-C3 and holo-C4, a pocket adjacent to the active site was also included into the binding site, identified by the authors of the MD study as a promising site for allosteric modulators.^22^ Validation of the docking protocol was performed on the only structure of a non-covalent complex, with inhibitor UK-371804. As such, the capability of the docking program to reproduce accurately the experimental complex was assessed by redocking UK-371804 on TMP2-NC with OpenEye FRED.^41^ The resulting Root Mean Square Deviation (RMSD) was of 1.97 Å, which is within the generally accepted threshold of 2.00 Å.^42,43^

**Fig. 3.**
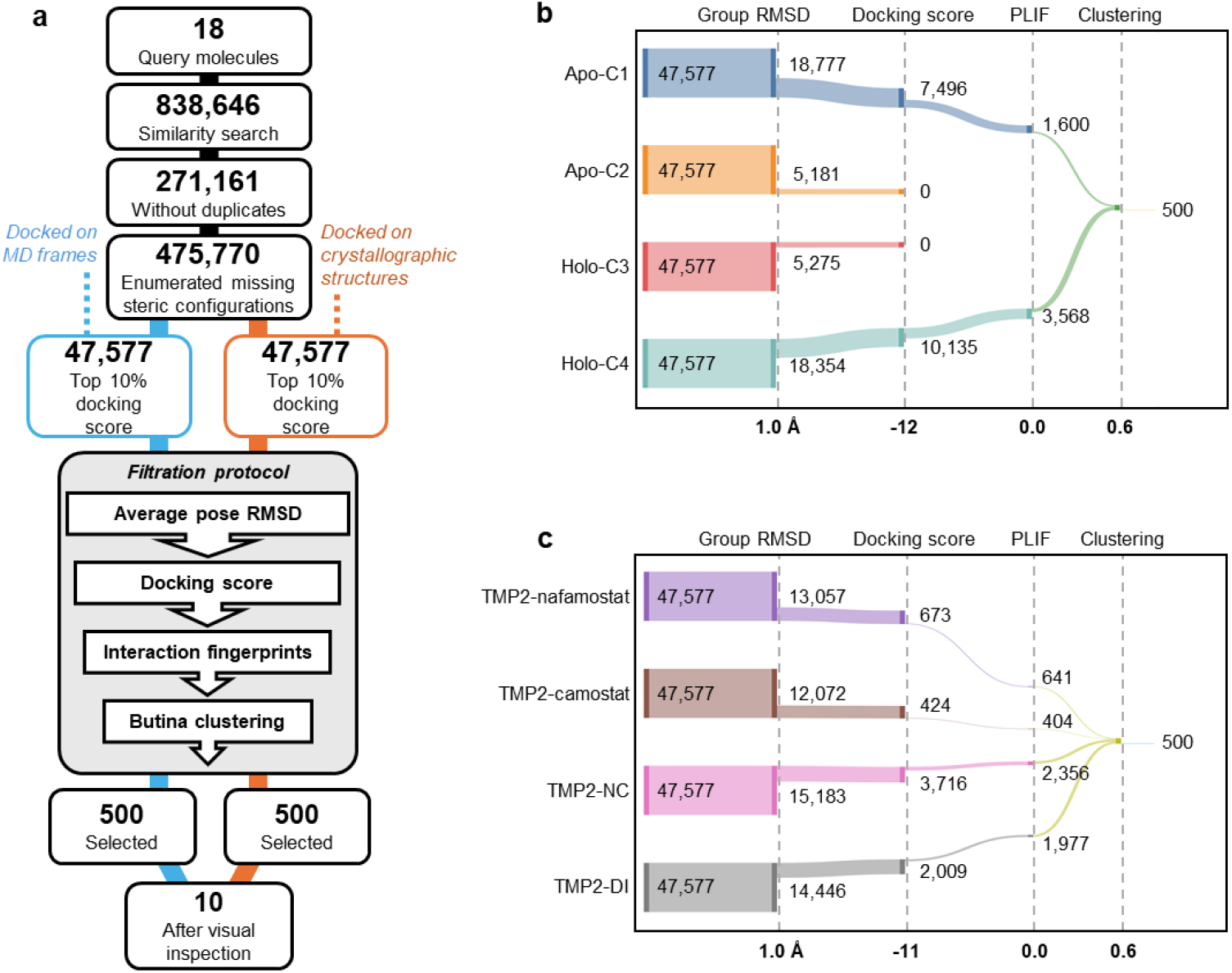
(a) General scheme of the filtration workflow. Light blue boxes and flow lines indicate poses generated on the MD frames, while orange ones are relative to X-ray crystallographic structures. The grey box represents the steps of the filtration workflow. (b,c) Sankey plots of the number of poses retained at each step of the filtration workflow for simulated and X-ray crystallographic target structures, respectively. Cutoff values for each filtration metric are indicated on the lower border of each graph. Between the two sets of structures they are the same except for docking score, with a cutoff of -12 for simulated structures and -11 for X-ray crystallographic structures.

Conformational states were successfully generated for 475,770 molecules of the previously prepared library, which were then docked onto each of the 8 protein structures using FRED. All molecules were successfully docked. From here on, we treated the docking results from the simulated structures and X-ray crystallographic structures separately. For each set of four structures, the best 10% of the library was selected. Since every ligand had four docking scores associated with it, the maximum rule was selected as consensus criterion, which considers only the best score among the four. This resulted in two different subsets of 47,577 compounds each, which were again subjected to conformer generation and molecular docking onto the 8 receptors, this time with “high resolution” settings. In this second run, 10 different poses were saved for each molecule. As such, the results were two sets of 47,577 compounds, each docked to 4 protein structures, meaning 40 scores for each compound and 1,903,080 total poses for each best-scoring subset.

The evaluation of such a large number of poses proved to be challenging. To tackle this problem and mitigate the risks associated with visual inspection and the sole reliance on docking scores, we introduced supplementary criteria to identify the most promising compounds. To this end, we developed an automatized and reproducible Python workflow which sequentially filters lists of docked compounds based on four metrics routinely applied and tested in the scientific community: pose coherence (i.e., the RMSD across the best 10 poses predicted for each compound), molecular docking score, interactions with key protein residues modelled with interaction fingerprints,^44^ and finally a cluster analysis to maximize the diversity of the final selection.^45^

An RMSD cutoff of 1.0 Å was selected as pose coherence cutoff, based on the RMSD distributions of docked compounds, which had a consistent peak around 0.7 Å across all structures. Compounds docked on apo-C2 and holo-C3 exhibited significantly worse RMSD distributions compared to all other structures, which showed consistent profiles with peaks around 0.6–0.7 Å. As a result, apo-C2 and holo-C3 were immediately penalized, retaining only about 5,000 compounds each, while apo-C1 and holo-C4 retained approximately 18,000 compounds each (**Fig. 3b**). The X-ray crystallographic structures yielded more uniform results, with the number of retained compounds ranging from 12,072 for TMP2-camostat to 15,183 for TMP2-NC (**Fig. 3c**).

The second filtration step applies a docking score threshold. Among the simulated structures, apo-C1 and holo-C4 displayed favorable score distributions with well-defined peaks around –12 and several values below –15, whereas apo-C2 and holo-C3 showed scores predominantly above –10. X-ray crystallographic structures had more uniform profiles, all peaking around –10, slightly worse than apo-C1 and holo-C4. Given the differences in scoring behavior, two score thresholds were applied: –12 for simulated structures and –11 for X-ray crystallographic ones. This resulted in the retention of 7,496 compounds for apo-C1 and 10,135 for holo-C4, while no compounds from apo-C2 and holo-C3 passed this filter. Among the X-ray crystallographic structures, TMP2-nafamostat and TMP2-camostat were the most penalized, with only 673 and 424 compounds retained, respectively, compared to 3,716 for TMP2-NC and 2,009 for TMP2-DI.

The third filtration step employed protein–ligand interaction fingerprints (PLIFs), using TMPRSS2 residues known to be important for substrate recognition and catalytic activity — namely His296, Asp345, Asp435, Gln438, Gly439, and Ser441 — as references. Only compounds showing at least one interaction with these key residues were retained. For apo-C1, only 1,600 molecules passed, compared with the 3,568 of holo-C4. Only 641 molecules passed for TMP2-nafamostat, 404 for TMP2-camostat, while TMP2-NC and TMP2-DI performed better, with 3,716 and 2,009 retained molecules, respectively.

For the final filtration step, clustering analysis was used to maximize chemical diversity among the candidates. Representative centroids were selected for visual inspection from the 500 most populated clusters, representing 90% of the total clustered compounds for the set of simulated receptors, and 88% for the set of X-ray crystallographic receptors. Visual inspection was conducted on both filtered sets of 500 compounds, as well as the original 18 query molecules. The 18 initial compounds were not automatically filtered but were instead manually evaluated based on the same metrics. Results for apo-C2, holo-C3, and all X-ray crystallographic structures were poor for these 18 molecules, so only the poses docked to apo-C1 and holo-C4 were further examined. For the two sets of 500 filtered compounds, one set consisted of molecules docked exclusively to apo-C1 (171 compounds) and holo-C4 (329), while the second set included molecules docked to TMP2-nafamostat (30), TMP2-camostat (34), TMP2-NC (174), and TMP2-DI (262).

Regarding visual inspection, the main criteria for the evaluation of the binding poses were steric complementarity with the binding site, followed by how many hydrogen bonds were made by the ligand and with which residues. Interactions with the residues used in the interaction fingerprint filtration step were valued highly, especially with Asp435, involved in recognition and correct orientation of TMPRSS2 substrates, and Ser441, the core residue of the catalytic process. An important criterion for exclusion was the presence of solvent-exposed hydrophobic groups, and other electrostatic incompatibilities, e.g., two adjacent negatively-charged groups, a hydrophilic moiety oriented towards a hydrophobic patch of the protein, or *vice versa*.

A set of 10 final compounds were selected for a first round of experimental testing (**Fig. 4a**). In order to identify the compounds, they were given a number relative to their place in one of the two best-scoring 10% subsets, preceded by “S-” for the simulated structures, and “C-” for the X-ray crystallographic structures. Among the 18 initial compounds, we selected molecules **6** (score -13.02) and **7** (score -14.60) for further studies, despite **6** having a group RMSD of 1.39 Å which would have theoretically deemed its exclusion during the filtration workflow, because starting compounds were of particularly high interest since already available for wet lab testing.

**Fig. 4.**
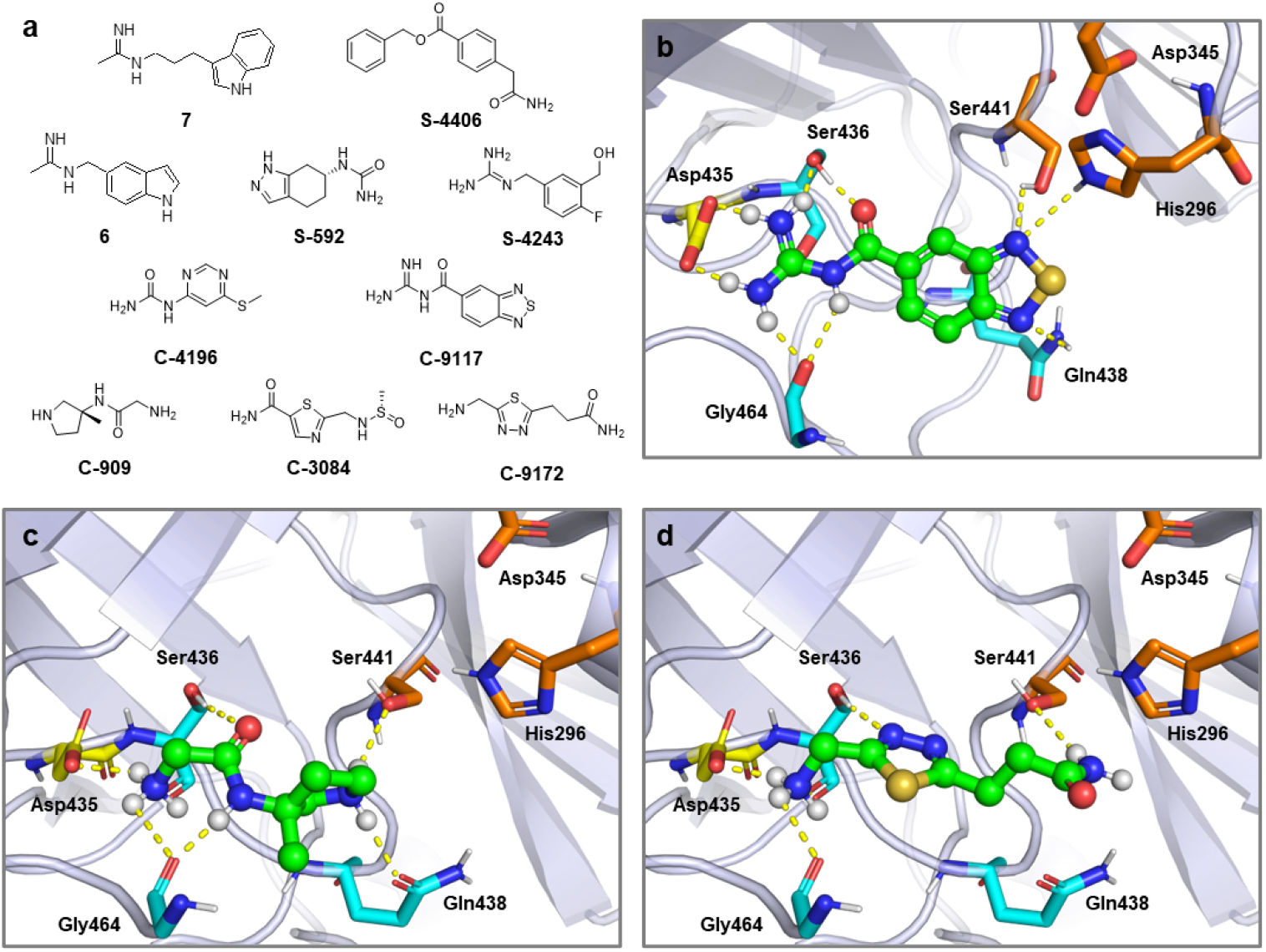
(a) Chemical structures of the 10 hit candidates selected from visual inspection. Compounds belonging to the 18 initial molecules are identified by a single number, compounds selected for a pose on one of the simulated structures are identified by “S” before a number corresponding to their position among the best 10% subset, compounds selected for a pose on one of the X-ray crystallographic structures are identified by “C” before the corresponding number. (b) Binding pose of hit candidate C-9117, docked on TMP2-NC. (c) Binding pose of hit candidate C-909, docked on TMP2-DI. (d) Binding pose of hit candidate C-9172, docked on TMP2-DI. All ligands are shown in ball-and-stick representation with green carbons, while key residues of the catalytic site are shown in sticks. The catalytic triad is shown with orange carbons, Asp435 in yellow, the other interacting residues Ser436, Gly439, Gly464 in light blue. Other non-hydrogen atoms are colored according to the standard color scheme. Interactions between the ligand and the residues are shown as yellow dashed lines.

In order to validate the filtration workflow, a set of 16 known TMPRSS2 inhibitors exhibiting *in vitro* or *in vivo* activity, including those co-crystallized with the target structures of this virtual screening, was sourced from the literature,^23,46–48^ their structures were prepared and subjected to high resolution docking under the same conditions as the main library. The results were passed through the filtration workflow, excluding the final clustering. The most penalising stage was the docking score cutoff, whereas the other metrics often yielded good values and were used with the same cutoff values as the main library. This resulted in 10 inhibitors being retained at the end by applying a score cutoff of –9 (NCGC00241049, NCGC00386945, NCGC00387094, NCGC00485967, NCGC00634982, gabexate, nafamostat, otamixaban, UK-371804, diminazene), 8 inhibitors retained for a –10 score cutoff (NCGC00241049, NCGC00386945, NCGC00634982, gabexate, nafamostat, otamixaban, UK-371804, diminazene), and only 3 passing with a docking score cutoff of –11 (NCGC00386945, NCGC00634982, nafamostat).

## Discussion

In this work, a virtual screening campaign was conducted to identify novel non-covalent inhibitors of TMPRSS2, a protease of clinical relevance due to its roles in prostate cancer and SARS-CoV-2 infection.

Starting from a curated set of 18 iNOS inhibitors bearing ionizable nitrogen-rich groups, a chemical library of 475,770 small molecules was generated. Eight TMPRSS2 receptor configurations were employed — four from MD simulations and four from recently solved X-ray crystal structures — to account for conformational diversity and pocket flexibility. Molecular docking was performed for the whole library across all structures, using FRED, and results were filtered through a deterministic, four-step pipeline that included RMSD analysis, docking score thresholds, interaction fingerprint screening, and clustering for chemical diversity. This process yielded two final sets of 500 compounds, one per structural group, which were then subjected to visual inspection along with the original 18 query compounds. Ten candidates were ultimately selected as promising hits.

The screening strategy was designed to maximize structural diversity, both in terms of ligands and receptor conformations. Receptor selection proved critical: while apo-C1 and holo-C4 (simulated structures) performed well, apo-C2 and holo-C3 failed to generate viable poses, emphasizing the impact of initial protein conformation on docking outcomes. Visual inspection of binding poses highlighted that the binding pockets of apo-C2 and holo-C3 were noticeably shallower than apo-C1 and holo-C4, respectively; this prevented ligands from making stable interactions and explains why virtual screening on these two structures was effectively a failure. In particular, the highest scoring poses among all structures were generated for the allosteric pocket of holo-C4, which is bordered on the bottom by the key residue Asp435, highlighting the potential for allosteric inhibition of TMPRSS2 through interaction with this substrate recognition residue. Further studies are needed to verify the stability of this pocket and its viability as a binding site.

Visual inspection allowed us to identify 10 hit candidates with the most promising binding modes and interaction patterns. For apo-C1, query molecule **7** and compound **S-4406** were selected. Both molecules orient a hydrophobic aromatic ring towards the hydrophobic bottom of the pocket. On the opposite end, for **7** a positively-charged amidine moiety makes hydrogen bonds with Asp458, Gly442 and Asp440, whereas in **S-4406** a terminal amide makes two hydrogen bonds with Gly282 and the backbone oxygen of catalytic Ser441.

For holo-C4, query molecule **6**, and compounds **S-592** and **S-4243** were selected. Compounds **6** and **S-4243** make a hydrogen bond with Asp435 with an amidine and guanidine group respectively, both interacting also with Thr459, and **S-4243** also makes a hydrogen bond with Ser382. On the outer side, they interact with Gly258 and Asn398, respectively. Compound **S-592** ureido group makes hydrogen bonds with Ala386, Asn398 and Asp440 on the outer side, while its tetrahydro-indazole moiety interacts with Asp435 backbone.

For the X-ray crystallographic structures, **C-4196** and **C-9117** were selected for TMP2-NC, and **C-909, C-3084**, and **C-9172** for TMP2-DI. **C-4196, C-9117** and **C-3084** share a similar scaffold, with a terminal polar moiety, either a urea, a guanidine or an amide, connected to an aromatic system, and various electronegative atoms on the opposite side. **C-4196** and **C-3084** instead share two amino/amide groups at the opposite sides of their structures, and an electron-rich center portion, corresponding respectively to an amide or an heterocyclic aromatic ring. All their interaction profiles are fairly similar. All five compounds interact with Asp435, and make hydrogen bonds with residues Ser436 and Gly464 of the S1 subsite, and with catalytic Ser441. **C-9117** also makes an additional hydrogen bond with His296. The positively-charged guanidine group of **C-9117** and linear amines of **C-909** and **C-9172** are able to make a salt bridge with Asp435, making their binding modes particularly akin to those of the known inhibitors co-crystallized in the original experimental complexes, where a salt bridge between an ionized guanidine group and Asp435 correctly orients the ligand to allow interaction with Ser441. Thus, these last three ligands were deemed particularly worthy of interest.

This study introduces a flexible, open-source computational workflow that integrates structure-based metrics and ligand-protein interaction profiles into an automatized filtering strategy. Its strengths lie in its automation, the inclusion of multiple target conformations and pockets, and, most importantly, reproducibility, with the final aim of mitigating some accuracy and subjectivity issues of visual inspection. Moreover, all compound prioritizations were based on metrics commonly accepted in the computational chemistry community. The code for this filtration pipeline is publicly available on GitHub at the following repository: https://github.com/LBIC-biocomp/filterfiesta/.

Nevertheless, limitations remain. Like most studies, the docking protocol was constrained to rigid receptor structures; moreover, the selected compounds are yet to undergo *in vitro* validation, and the final selection still required a degree of subjective visual inspection. Furthermore, only non-covalent interactions were explored, leaving out the potential of covalent inhibition strategies.

Future works will focus on experimental validation of the top-ranked candidates, and the integration of protein flexibility during docking or rescoring stages, with the goal of improving predictive accuracy and advancing these hits toward early-stage drug development.

## Methods

### Library design and preparation

The compound library utilised in this project is a subset of Enamine REAL database, where REAL stands for REadily AccessibLe.^40^ It is a database which, as of February 2025, comprises over 9.6 billion molecules, all designed to comply with Lipinski rule of five and Veber criteria.^40^ To select a suitable subset of the REAL database, Smallworld search engine – by NextMove Software^39^ – was used to search for compounds similar to query molecules. Smallworld represents molecules in terms of graph theory, in which a molecule’s atoms are symbolised by objects called vertices or nodes, visualised as points, which are connected by edges, representing chemical bonds. This allows to define the similarity between two molecules using the concept of Graph Edit Distance (GED), i.e., the minimum number of insertions, deletions and substitutions of nodes and edges required to turn one graph into the other.^49^ The default search parameters were used for the similarity search, corresponding to a GED cutoff of 12. Within the OpenEye workflow, protonation states were calculated at a pH of 7.4 with the tool FixpKa, and stereoisomer generation was managed with Flipper for molecules with unassigned stereochemistry, with enumeration of both R/S and E/Z all possible configurations. The OpenEye dedicated program OMEGA was used to convert the 2D structures into an ensemble of conformers. Up to 200 conformers per ligand are calculated under default parameters (“classic” mode), a number which goes up to 800 for its high-resolution mode (“pose” mode).

### Molecular docking

In this virtual screening, ensemble docking was performed on two separate sets of receptors, each composed of four structures of TMPRSS2 sourced from the literature. The first four structures are MD frames obtained from the work of Frumenzio *et al*.^22^ Briefly, the authors started from the X-ray crystallographic structure of TMPRSS2 in complex with nafamostat (PDB ID 7MEQ), with the transmembrane helix and LDLRA domain modelled with AlphaFold2. The four configurations were identified by the authors as the most representative from 2 µs-long trajectories of the apo and holo forms of the protein. The second set of structures came from the work of Wang *et al*.,^23^ who published the most complete X-ray crystallographic structures of TMPRSS2 available as of September 2024. The four structures each depict the protease in complex with a different inhibitor, and are TMPRSS2 in complex with nafamostat (PDB ID 7XYD); in complex with camostat (PDB ID 7Y0E); in complex with non-covalent inhibitor UK-371804 (PDB ID 7Y0F); in complex with nafamostat-inspired dual inhibitor 212-148 (PDB ID 8HD8).

The four X-ray crystallographic structures were originally saved as dimers, so their PDB files were opened in ChimeraX and the excess monomers removed. Furthermore, for structures in complex with covalent inhibitors (i.e., holo-C3, holo-C4, TMP2-nafamostat, TMP2-camostat, TMP2-DI), the respective covalent ligands were removed and the hydroxyl group of Ser441 was rebuilt. Next, the structures were processed with the MakeReceptor software and boxes delimiting the active site were generated centred on Ser441, selecting the “molecular” option for calculations of the active site volume. A binding site of bigger volume (2141 Å^3^) was delineated for holo-C4 and holo-C3 in order to better encompass the putative allosteric pocket, whereas the other binding site volumes were relatively similar, ranging from 1540 to 1770 Å^3^. OpenEye FRED was used to dock the prepared library onto the receptors, using the default resolution for the whole library, and the “high resolution” mode for the top 10% best scoring molecules. For the high resolution docking, 10 poses were saved for each molecule.

For the redocking of UK-371804 onto TMP2-NC, the inhibitor in SMILES notation was processed by the program OMEGA for the generation of conformers. Then, the conformers were docked with FRED on TMP2-NC using “high resolution” mode. The resulting docked structure, stored in PDB format, was opened in the program ChimeraX together with the experimental complex, and a built-in function of the program was used to measure the RMSD between the two ligand structures.

### Molecular similarity

We used 1024-bits Extended Connectivity Fingerprints (ECFPs) and the Tanimoto index to quantify the similarity between molecules. The similarity index works by comparing common features or bits (for continuous and binary variables respectively) relative to the total number of features/bits. The result is a number which can take any value between 0 and 1, with 1 meaning the two compared molecules are identical. The fingerprints and similarities were calculated via the RDKit v. 2024.3.5 GetMorganGenerator function with radius=4 and BulkTanimotoSimilarity, respectively.

### Binding poses coherence

The coherence of the binding modes was assessed by considering multiple poses for each ligand and calculating the average RMSD within the group of poses. Specifically, using the RDKit toolkit,^50^ each group of poses was converted into molecule objects and subsequently used to generate an average molecular structure. The RMSD between this average structure and each individual pose was then calculated using RDKit’s built-in functions. Finally, the mean RMSD across all poses was computed to assess the pose coherence of the molecule.

### Protein-Ligand Interaction Fingerprints

The Open Drug Discovery Toolkit (ODDT) Python library^51^ enables straightforward calculation of various types of protein–ligand interactions and stores the resulting data as numerical fingerprints. For each interaction between the ligand and a protein residue, the value corresponding to that residue and interaction type is incremented by one. A value of zero indicates the absence of that interaction type. Each fingerprint encodes eight types of interactions: hydrophobic contacts; aromatic face-to-face and edge-to-face interactions; hydrogen bonds, where the protein acts as either donor or acceptor; salt bridges involving positively or negatively charged protein residues; and salt bridges formed through ionic bonds with metal ions.

This was used to select compounds with interaction profiles complying with a chosen template. First, all interaction fingerprints were converted into binary vectors, by forcing all non-zero values to 1. Then, an analogous binary interaction fingerprint was created as reference, containing all the desired interactions with values set to 1. Finally, the Jaccard similarity to the reference was calculated for each interaction fingerprint. Compounds with an acceptable interaction profile were defined as those with a non-zero Jaccard value, meaning at least one interaction in common with the reference.

### Clustering

Cluster analysis was performed using the Butina algorithm,^45^ which utilises a Jaccard similarity cutoff as the only input. The algorithm identifies the molecule fingerprint with the highest number of neighbours (i.e., molecules fingerprints within a certain similarity value) which is taken as cluster centroid. All the neighbouring molecules within the selected similarity threshold are assigned to the new cluster and removed from the list. The process is repeated until all the list of molecules has been processed. In this work, a similarity cutoff of 0.6 was used.

## Acknowledgments

F.M. was supported by Ministero dell’Istruzione, dell’Università e della Ricerca (RFO grant 2024) and by Consorzio Interuniversitario Risonanze Magnetiche di Metallo Proteine (CIRMMP). OpenEye software was used under free academic licensing.

## Data Availability

The complete library in SMILES format and docking scores are available upon request. Interested parties are kindly invited to contact the corresponding authors to request access to the aforementioned data.

## References

1. Paoloni-Giacobino, A., Chen, H., Peitsch, M. C., Rossier, C. & Antonarakis, S. E. Cloning of the TMPRSS2 Gene, Which Encodes a Novel Serine Protease with Transmembrane, LDLRA, and SRCR Domains and Maps to 21q22.3. Genomics 44, 309–320 (1997).

2. Zheng, X. L., Kitamoto, Y. & Sadler, J. E. Enteropeptidase, a type II transmembrane serine protease. FBE 1, 242–249 (2009).

3. Fasquelle, L. et al. Tmprss3, a Transmembrane Serine Protease Deficient in Human DFNB8/10 Deafness, Is Critical for Cochlear Hair Cell Survival at the Onset of Hearing. J of Biol Chem 286, 17383–17397 (2011).

4. Li, S., Wang, L., Sun, S. & Wu, Q. Hepsin: a multifunctional transmembrane serine protease in pathobiology. FEBS J 288, 5252–5264 (2021).

5. Li, H., Zhang, Y. & Wu, Q. Role of corin in the regulation of blood pressure. Curr Opin Nephrol Hypertens 26, 67–73 (2017).

6. Fraser, B. J. et al. Structure and activity of human TMPRSS2 protease implicated in SARS-CoV-2 activation. Nat Chem Biol 18, 963–971 (2022).

7. Bugge, T. H., Antalis, T. M. & Wu, Q. Type II Transmembrane Serine Proteases. J Biol Chem 284, 23177–23181 (2009).

8. Vaarala, M. H., Porvari, K. S., Kellokumpu, S., Kyllönen, A. P. & Vihko, P. T. Expression of transmembrane serine protease TMPRSS2 in mouse and human tissues. J Pathol 193, 134–140 (2001).

9. Chen, Y.-W. et al. TMPRSS2, a Serine Protease Expressed in the Prostate on the Apical Surface of Luminal Epithelial Cells and Released into Semen in Prostasomes, Is Misregulated in Prostate Cancer Cells. Am J Pathol 176, 2986–2996 (2010).

10. Ko, C.-J. et al. Androgen-Induced TMPRSS2 Activates Matriptase and Promotes Extracellular Matrix Degradation, Prostate Cancer Cell Invasion, Tumor Growth, and Metastasis. Cancer Res 75, 2949–2960 (2015).

11. Gioukaki, C. et al. Unravelling the Role of P300 and TMPRSS2 in Prostate Cancer: A Literature Review. Int J Mol Sci 24, 11299 (2023).

12. Wettstein, L., Kirchhoff, F. & Münch, J. The Transmembrane Protease TMPRSS2 as a Therapeutic Target for COVID-19 Treatment. Int J Mol Sci 23, 1351 (2022).

13. Matsuyama, S. et al. Efficient Activation of the Severe Acute Respiratory Syndrome Coronavirus Spike Protein by the Transmembrane Protease TMPRSS2. J Virol 84, 12658–12664 (2010).

14. Kleine-Weber, H., Elzayat, M. T., Hoffmann, M. & Pöhlmann, S. Functional analysis of potential cleavage sites in the MERS-coronavirus spike protein. Sci Rep 8, 16597 (2018).

15. Bertram, S. et al. TMPRSS2 Activates the Human Coronavirus 229E for Cathepsin-Independent Host Cell Entry and Is Expressed in Viral Target Cells in the Respiratory Epithelium. J Virol 87, 6150–6160 (2013).

16. Shirato, K., Kawase, M. & Matsuyama, S. Wild-type human coronaviruses prefer cell-surface TMPRSS2 to endosomal cathepsins for cell entry. Virology 517, 9–15 (2018).

17. Milewska, A. et al. Entry of Human Coronavirus NL63 into the Cell. J Virol 92, 10.1128/jvi.01933-17 (2018).

18. Saunders, N. et al. TMPRSS2 is a functional receptor for human coronavirus HKU1. Nature 624, 207–214 (2023).

19. Schechter, I. & Berger, A. On the size of the active site in proteases. I. Papain. Biochem Biophys Res Commun27, 157–162 (1967).

20. Hooper, J. D., Clements, J. A., Quigley, J. P. & Antalis, T. M. Type II Transmembrane Serine Proteases: Insights into an Emerging Class of Cell Surface Proteolytic Enzymes. J Biol Chem 276, 857–860 (2001).

21. Hedstrom, L. Serine Protease Mechanism and Specificity. Chem Rev 102, 4501–4524 (2002).

22. Frumenzio, G. et al. Conformational response to ligand binding of TMPRSS2, a protease involved in SARS-CoV-2 infection: Insights through computational modeling. Proteins 91, 1288–1297 (2023).

23. Wang, H. et al. Structure-based discovery of dual pathway inhibitors for SARS-CoV-2 entry. Nat Commun 14, 7574 (2023).

24. Drag, M. & Salvesen, G. S. Emerging principles in protease-based drug discovery. Nat Rev Drug Discov 9, 690–701 (2010).

25. Mantzourani, C., Vasilakaki, S., Gerogianni, V.-E. & Kokotos, G. The discovery and development of transmembrane serine protease 2 (TMPRSS2) inhibitors as candidate drugs for the treatment of COVID-19. Expert Opin Drug Discov 1–16 doi:10.1080/17460441.2022.2029843.

26. Hoffmann, M. et al. Nafamostat Mesylate Blocks Activation of SARS-CoV-2: New Treatment Option for COVID-19. Antimicrob Agents Chemother 64, e00754–20 (2020).

27. Khan, U. et al. Safety and Efficacy of Camostat Mesylate for Covid-19: a systematic review and Meta-analysis of Randomized controlled trials. BMC Infect Dis 24, 709 (2024).

28. Hernández-Mitre, M. P. et al. TMPRSS2 inhibitors for the treatment of COVID-19 in adults: a systematic review and meta-analysis of randomized clinical trials of nafamostat and camostat mesylate. Clin Microbiol Infect 30, 743–754 (2024).

29. Paggi, J. M., Pandit, A. & Dror, R. O. The Art and Science of Molecular Docking. Annu Rev Biochem 93, 389–410 (2024).

30. Fischer, A., Smieško, M., Sellner, M. & Lill, M. A. Decision Making in Structure-Based Drug Discovery: Visual Inspection of Docking Results. J Med Chem 64, 2489–2500 (2021).

31. Sadybekov, A. V. & Katritch, V. Computational approaches streamlining drug discovery. Nature 616, 673–685 (2023).

32. Xue, Q., Yan, Y., Zhang, R. & Xiong, H. Regulation of iNOS on Immune Cells and Its Role in Diseases. Int J Mol Sci 19, 3805 (2018).

33. Calabrese, V. et al. Nitric oxide in the central nervous system: neuroprotection versus neurotoxicity. Nat Rev Neurosci 8, 766–775 (2007).

34. Stern, A. M. & Zhu, J. Chapter Five - An Introduction to Nitric Oxide Sensing and Response in Bacteria. in Adv Appl Microbiol (eds. Sariaslani, S. & Gadd, G. M.) vol. 87 187–220 (Academic Press, 2014).

35. Yang, L. et al. The signal pathways and treatment of cytokine storm in COVID-19. Sig Transduct Target Ther 6, 255 (2021).

36. Carrión, M. D. et al. New amidine-benzenesulfonamides as iNOS inhibitors for the therapy of the triple negative breast cancer. Eur J Med Chem 248, 115112 (2023).

37. Maccallini, C. et al. Discovery of N-{3-[(ethanimidoylamino)methyl]benzyl}-l-prolinamide dihydrochloride: A new potent and selective inhibitor of the inducible nitric oxide synthase as a promising agent for the therapy of malignant glioma. Eur J Med Chem 152, 53–64 (2018).

38. Re, N., Fantacuzzi, M., Maccallini, C., Paciotti, R. & Amoroso, R. Recent Developments of Amidine-like Compounds as Selective NOS Inhibitors. Curr Enzyme Inhib 12, 30–39 (2016).

39. NextMove Software | SmallWorld. https://www.nextmovesoftware.com/smallworld.html.

40. REAL Database - Enamine. https://enamine.net/compound-collections/real-compounds/real-database.

41. OEDOCKING. OpenEye, Cadence Molecular Sciences. Santa Fe, NM. http://www.eyesopen.com.

42. Gohlke, H., Hendlich, M. & Klebe, G. Knowledge-based scoring function to predict protein-ligand interactions. J Mol Biol 295, 337–356 (2000).

43. Jiang, H. et al. Guiding Conventional Protein–Ligand Docking Software with Convolutional Neural Networks. J Chem Inf Model 60, 4594–4602 (2020).

44. Li, Z., Huang, R., Xia, M., Patterson, T. A. & Hong, H. Fingerprinting Interactions between Proteins and Ligands for Facilitating Machine Learning in Drug Discovery. Biomolecules 14, 72 (2024).

45. Butina, D. Unsupervised Data Base Clustering Based on Daylight’s Fingerprint and Tanimoto Similarity: A Fast and Automated Way To Cluster Small and Large Data Sets. J Chem Inf Comput Sci 39, 747–750 (1999).

46. Hu, X. et al. Discovery of TMPRSS2 Inhibitors from Virtual Screening as a Potential Treatment of COVID-19. ACS Pharmacol Transl Sci 4, 1124–1135 (2021).

47. Shapira, T. et al. A TMPRSS2 inhibitor acts as a pan-SARS-CoV-2 prophylactic and therapeutic. Nature 605, 340–348 (2022).

48. Xu, Y.-M., Inacio, M. C., Liu, M. X. & Gunatilaka, A. A. L. Discovery of diminazene as a dual inhibitor of SARS-CoV-2 human host proteases TMPRSS2 and furin using cell-based assays. Curr Res Chem Biol 2, 100023 (2022).

49. Sanfeliu, A. & Fu, K.-S. A distance measure between attributed relational graphs for pattern recognition. IEEE Trans Syst Man Cybern SMC-13, 353–362 (1983).

50. RDKit: Open-source cheminformatics. https://www.rdkit.org.

51. Wójcikowski, M., Zielenkiewicz, P. & Siedlecki, P. Open Drug Discovery Toolkit (ODDT): a new open-source player in the drug discovery field. J Cheminform 7, 26 (2015).

